# Observation of the Transport and Removal of Lipofuscin from the Mouse Myocardium using Transmission Electron Microscope

**DOI:** 10.1101/2020.03.10.985507

**Authors:** Lei Wang, Chang-Yi Xiao, Jia-Hua Li, Gui-Cheng Tang, Shuo-Shuang Xiao

## Abstract

This study was performed to investigate whether the lipofuscin formed within cardiomyocytes can be excluded by the myocardial tissue. We have provided indicators that can be used for future studies on anti-aging interventions.

In the present study, the heart of a 5-month-old BALB/c mouse was obtained for resin embedding and ultra-thin sectioning. The specimens were observed under a Hitach 7500 transmission electron microscope, and the images were acquired using an XR401 side-insertion device.

Lipofuscin granules are found abundantly in myocardial cells. Cardiomyocytes can excrete lipofuscin granules into the myocardial interstitium using capsule-like protrusions that are formed on the sarcolemma. These granules enter the myocardial interstitium and can be de-aggregated to form “membrane-like garbage”, which can pass from the myocardial stroma into the lumen of the vessel through its walls in the form of soluble fine particles through diffusion or endocytosis of capillaries. Smaller lipofuscin granules can pass through the walls of the vessels and enter the blood vessel lumen through the active transport function of the capillary endothelial cells. When the extended cytoplasmic end of macrophages and fibroblasts fuse with the endothelial cells, the lipofuscin granules or clumps found in the cells of the myocardial interstitium are transported to the capillary walls, and then, they are released into the lumen of the blood vessel by the endothelial cells.

The myocardial tissues of mice have the ability to eliminate the lipofuscin produced in the cardiomyocytes into the myocardial blood circulation. Although there are several mechanisms through which the myocardial tissues release lipofuscin into the bloodstream, it is mainly carried out in the form of small, fine, soluble, continuous transport.

## Introduction

It is well known that one of most important research areas in *in vivo* anti-aging intervention is to discover objective evaluation indicators. The content of lipofuscin in organs is an important indicator for assessing the aging status. Lipofuscin is a yellowish-brown pigment composed of highly oxidized proteins, lipids, and metals. Lipofuscin is widely observed in post-mitotic cells, especially in cells with long life spans such as neurons and cardiomyocytes[1,2]. Lipofuscin is often known as the “age pigment” and is considered a hallmark of aging. This is not only because the amount of lipofuscin increases with age, showing an approximately linear dependence, but also, and more importantly, because the rate of lipofuscin accumulation correlates negatively with longevity[3–7].

Lipofuscin, or age pigment, represents an intra-lysosomal polymeric material that cannot be degraded by lysosomal hydrolases. Although its continuous accumulation over time in post-mitotic cells, such as neurons and cardiomyocytes, has been known for more than 100 years, it has recently been suggested that accumulated lipofuscin may be hazardous to cellular functions[6, 8, 9]. There are strong indications that progressive deposition of lipofuscin ultimately decreases cellular adaptability and promotes the development of age-related pathologies, including neuro-degenerative diseases, heart failure, and macular degeneration[8, 9]. von Zglinicki T *et al*. found that accelerated lipofuscin accumulation is sufficient to block cellular proliferation within a short period time and induce cell death. They concluded that accumulation of lipofuscin in post-mitotic cells is not just a harmless consequence of ageing, but rather one of the most important causes of senescence[10].

Some researchers believe that the accumulation of lipofuscin within post-mitotic cells occurs primarily because it is non-degradable and cannot be removed from the cells through exocytosis[11, 12].

However, previous studies have shown that lipofuscin might be released from tissues or cells. A study by El-Ghazzawi *et al*. strongly suggested that nerve cells physiologically remove lipofuscin into the capillary endothelium[13]. Upon treatment with meclofenoxate, there is a gradual decrease in the volume of the pigment in the myocardium. After a treatment period of 4-6 weeks, the pigment bodies were found to be lodged in the capillary endothelium and the lumen, which facilitates the removal of the pigment through the blood stream[14]. Joris *et al*. observed the formation and outcome of lipofuscin in human coronary endothelium, and showed that the bulging cell processes containing lipofuscin probably eventually break off and are shed into the bloodstream[15]. Similarly, it has also been proposed that neuronal lipofuscin was transported to the capillary wall and extruded into the lumen in the avian brain[16]. However, some researchers have questioned the studies showing the exclusion of lipofuscin from tissues[17].

Hence, it is still not conclusive whether the non-dividing cells in tissues and organs possess a mechanism for excluding lipofuscin. To ascertain whether the heart has a mechanism for eliminating lipofuscin from the myocardium, we carried out investigations using the mouse heart.

## 1. Materials and Methods

Five-month-old BALB/c mice were purchased from Laboratory Animal Center of our institution. All applicable international, national and institutional guidelines for the care and use of animals were followed. The animals were anesthetized using ether. The chest was opened and the heart was removed. Small tissue blocks were obtained from the anterior wall of the left ventricle and were fixed with 1.5% glutaraldehyde. The fixed tissue blocks were further treated with 1% osmium tetroxide, dehydrated, and embedded in epoxy resin. The specimens were sliced using the Microm HM340E rotary ultramicrotome produced by Leica Microsystems Heidelberg GmbH in Germany, and the ultra-thin sections were placed on a copper screen and stained again using lead nitrate and uranyl acetate. Finally, specimens were observed with a Hitachi 7500 transmission electron microscope, and images were acquired using an AMT XR401 side insertion drawing equipment produced by Advanced Microscopy Techniques in USA.

## 2. Results

### 2.1. Myocardial fibers (Cardiac cells)

Microscopic observation of myocardial tissue revealed clearly visible lipofuscin deposits that were scattered around the myocardial fibers (cardiac cells) and myocardial interstitium. These lipid-containing, brown deposits were granular or agglomerated; widely varied in size and form; displayed high or medium electron density; uniform or uneven internal structure; or lamellar or lipid droplets; with or without membrane coating; or dispersed or aggregated into large clumps.

Lipofuscin found in the myocardial fibers was mainly distributed in the cytoplasm-rich regions at both ends of the nucleus and was interspersed between a large numbers of mitochondria. The distribution of lipofuscin granules was lower in the myofibrils that were far from the nucleus (Fig 1). Lipofuscin granules could also be seen at the edges of myocardial fibers and below the sarcolemma (Fig 2, 15, 16).

**Fig 1.**
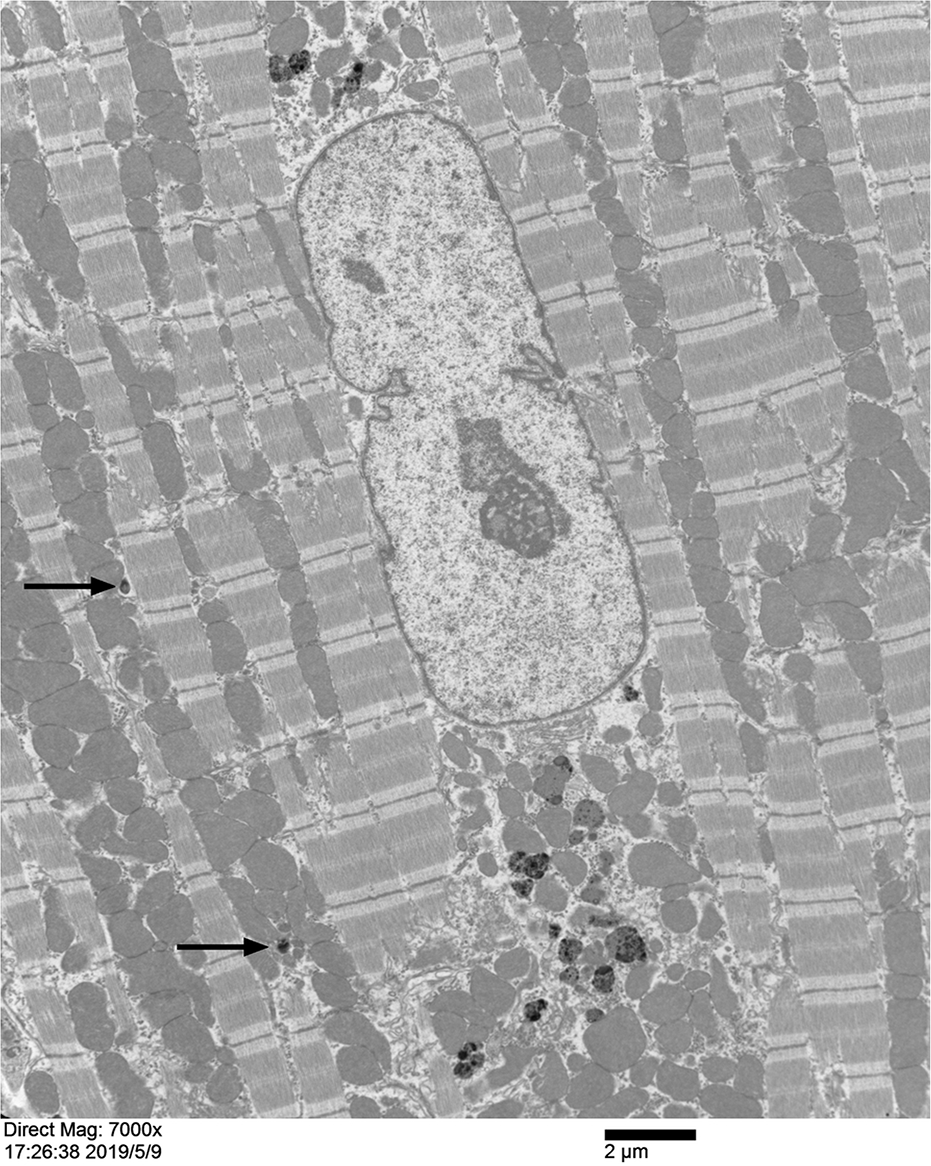
Longitudinal section of myocardial fiber in mouse heart. In the vertical axis of the myocardial fiber in the figure, an oval-shaped cell nucleus can be seen. A large number of densely arranged mitochondria can be seen in the cytoplasm at both ends of the nucleus, and there are multiple lipofuscin granules with high electron density among the mitochondria. These lipofuscin granules are of different sizes and uneven internal structures. Several scattered lipofuscin granules can also be seen among the myofibrils (shown by black arrows). (×15000, scale plate: 2 μm)

**Fig 2.**
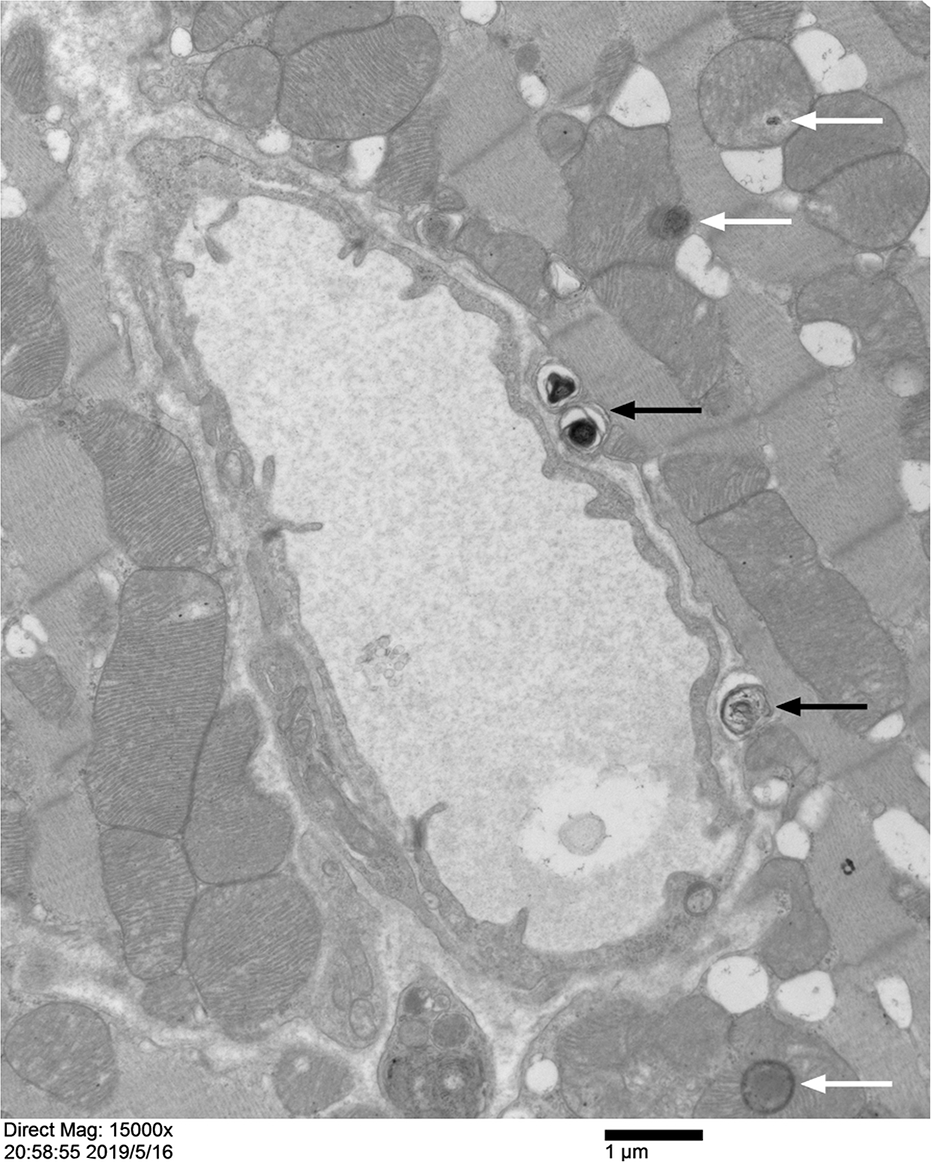
Myocardial fibers and capillary in mouse heart. Three lipofuscin granules with high electron density can be seen within the myocardial fiber in the right half of the picture close to the capillary wall. They protrude from the surface of the myocardial fiber covered only by a layer of sarcolemma (shown by black arrows). There is no basement membrane outside the capillary endothelium. In the deeper part of the myocardial fiber, two lipofuscin-like granules with different electron densities can be seen (shown by white arrows). (×15000, scale plate: 1 μm)

Compared with the lipofuscin granules located deep in the cardiomyocytes, the granules located around the cardiomyocytes were significantly smaller. The sarcolemma on the surface of cardiomyocytes was clearly visible, but often displayed unevenness. Bulging, rod-, sac-, or capsule-like protrusions of different sizes and lengths, were often observed, which contained the lipofuscin granules (Fig 3, 4, 9, 13).

**Fig 3.**
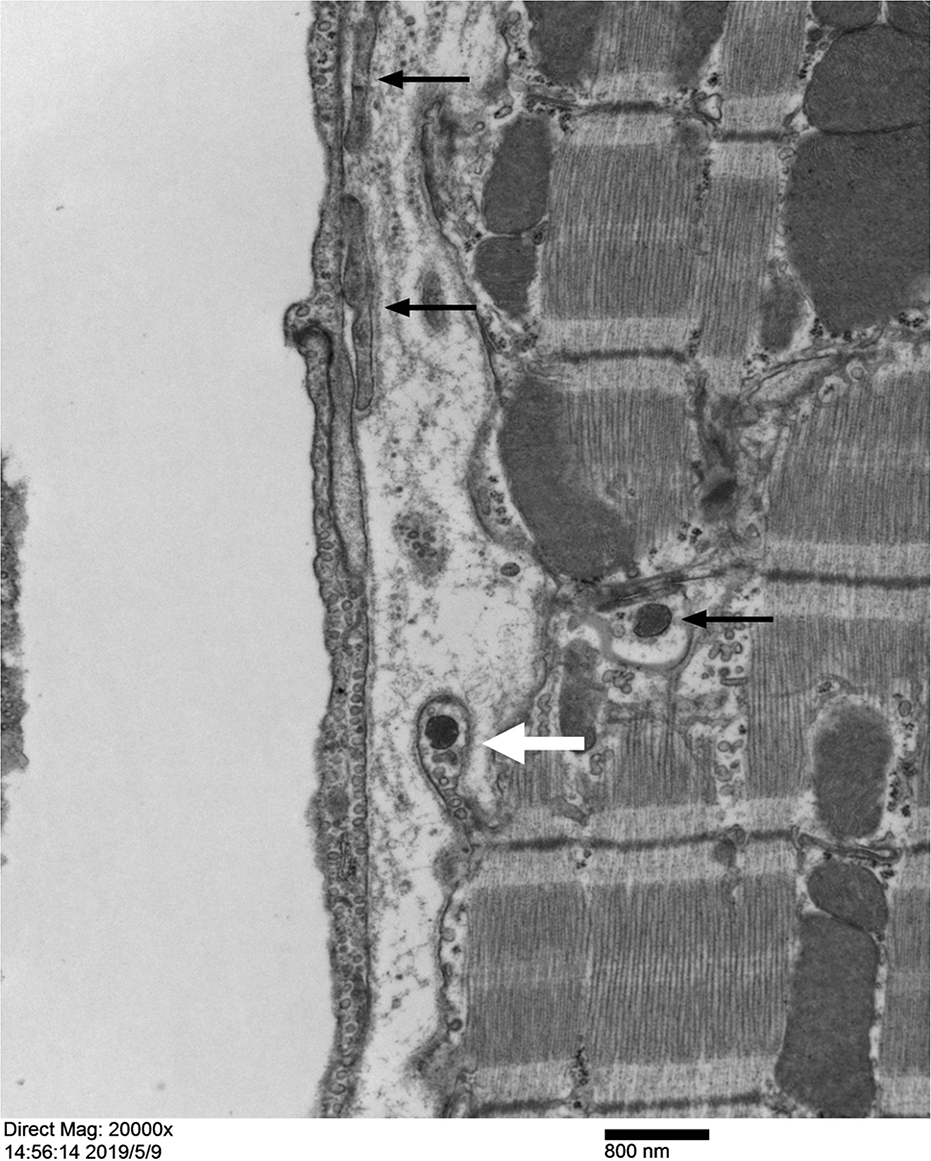
Myocardial fiber and capillary in mouse heart. The right-hand side of the figure shows a partial longitudinal cut of a myocardial fiber. A capsule-like protrusion can be seen on the surface of the myocardial fiber (shown by a white thick arrow). In the inner center of the top of the capsule-like protrusion, a high electron density spherical lipofuscin granule can be seen. A lipofuscin granule similar to that present in the protrusion can also be seen inside the myocardial fibers near the site where the capsule-like protrusion exits (shown by a black arrow). The capillary endothelium is continuous and intact, and its cytoplasm contains numerous vesicles. The basement membrane outside the endothelium is thin and discontinuous. There are two fibroblast terminal cytoplasmic fragments attached to the upper part outside the capillary wall (shown by white thin arrows). (×20000, scale plate: 800 nm)

**Fig 4.**
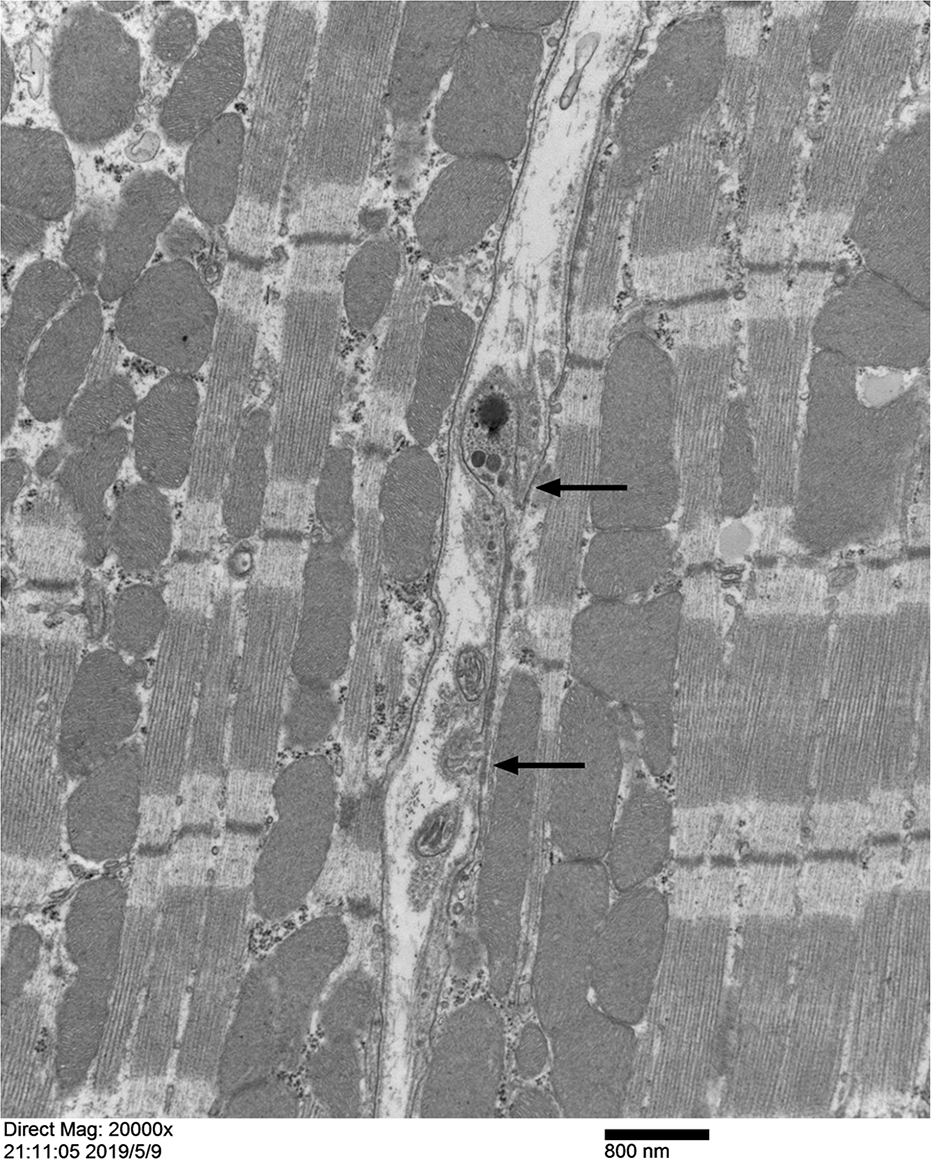
Myocardial fiber and myocardial interstitium in mouse heart. This image shows a longitudinal section of the myocardial fiber. A narrow gap can be seen between the two myocardial fibers. The sarcolemma on the surface of the myocardial fiber on the left side of the gap is clearly visible, and the walking is smooth. The surface of the myocardial fiber on the right side of the gap is uneven, and there are several capsule-like protrusions on the surface of the myocardial fiber (shown by black arrows) of different sizes, heights, and widths. In the middle region of the cytoplasm inside the larger protrusion, there are several lipofuscin granules with high electron density. (×20000, scale plate: 800 nm)

### 2.2. Cells in myocardial interstitium

In addition to being rich in capillaries, the interstitial space between myocardial fibers also had two types of cells, fibroblasts and macrophages, with fibroblasts being the predominant type.

Fibroblasts are slender spindle-shaped cells and their cell bodies extend along the longitudinal direction of the myocardial fibers with both their ends stretching far. The surface of fibroblast is smooth with no visible phagocytosis. Fibroblasts have a dense cytoplasmic structure and abundant organelles, which contain abundant rough endoplasmic reticulum, free ribosomes, and mitochondria (Fig 5, 6, 7, S1 Fig).

**Fig 5.**
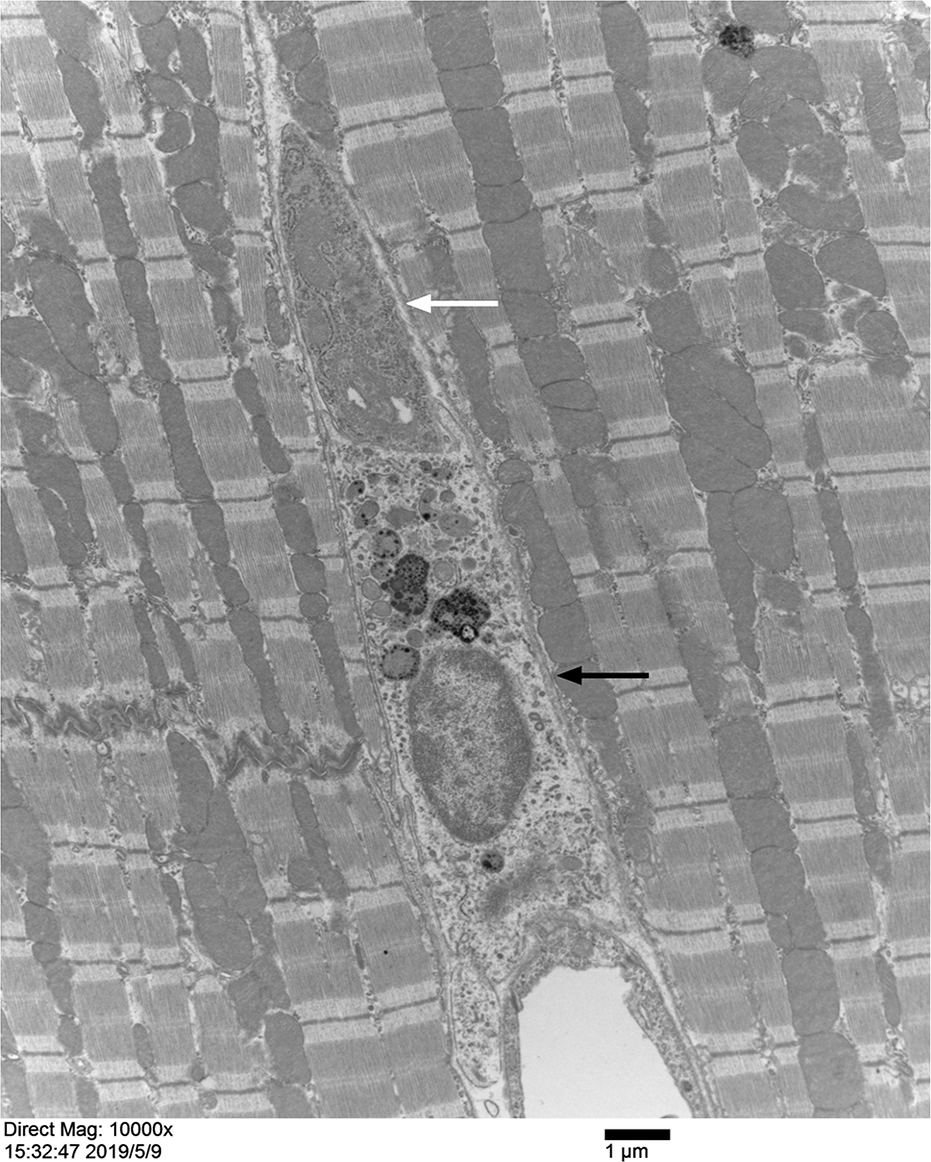
Myocardial interstitium of mouse heart. Arranged from the top to the bottom in the cardiac interstitium, a fibroblast (shown by a white arrow), a macrophage (shown by a black arrow), and capillary. The fibroblast has a high density of organelles in the cytoplasm, with abundant rough endoplasmic reticulum, free ribosomes, and mitochondria. The macrophage has relatively low cytoplasmic density, rich cytoplasm, abundant vesicles including phagocytic vesicles, and more lipid-like lipofuscin masses. These lipofuscin masses differ in size, internal structure, and electron density. The macrophage is closely attached to the capillariy below, and the basement membrane outside the capillary in this area is incomplete or even missing. (×10 000, scale plate: 1 μm)

**Fig 6.**
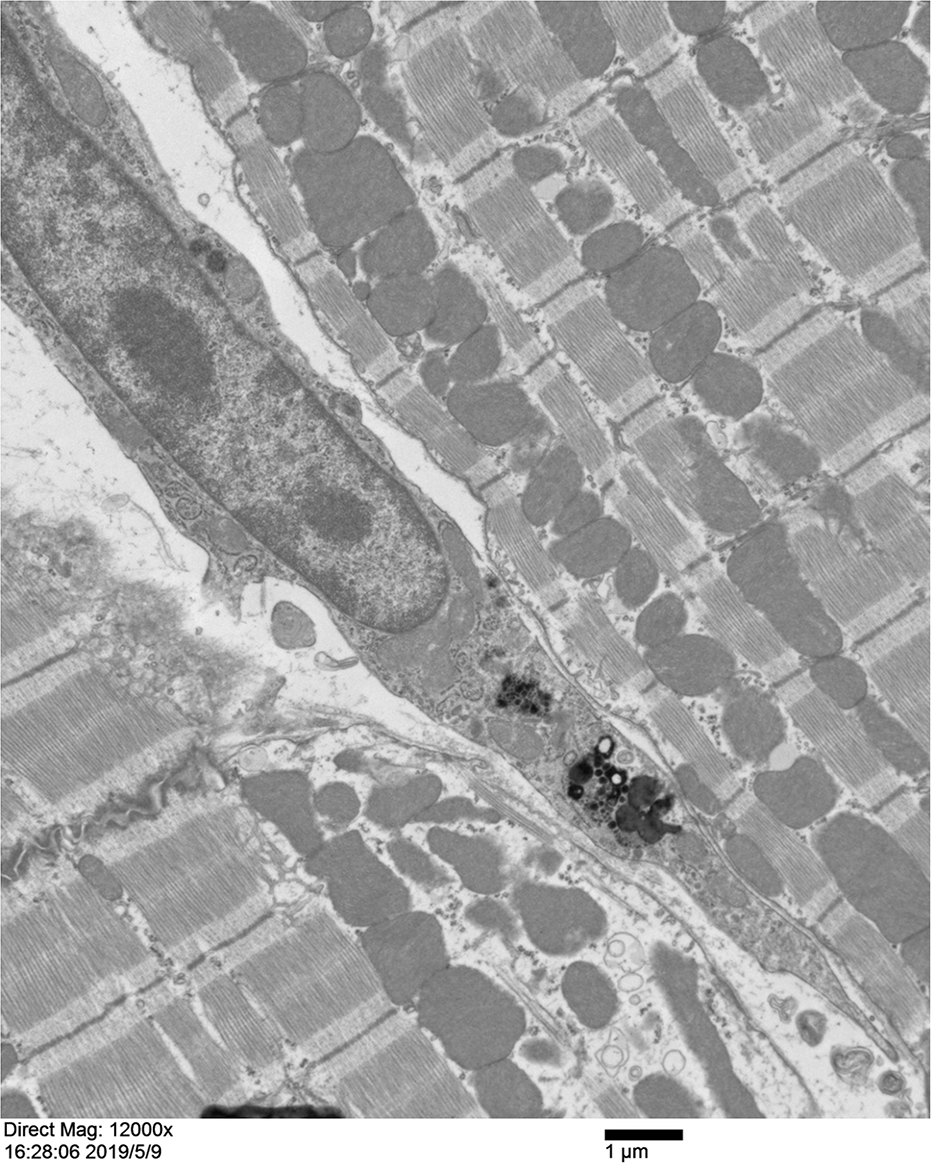
Fibroblast in myocardial interstitium of mouse heart. A fibroblast is long spindle-shaped cell that is present in the longitudinal direction of the myocardial fibers with slender ends. The cytoplasm has high density and is rich in organelles. It also contains lipofuscin granules or clumps of varying sizes. These lipofuscinous granules or clumps have uneven internal structures and different morphologies, have high electron density, and have no membrane coating on the periphery. (×12 000, scale plate: 1 μm)

**Fig 7.**
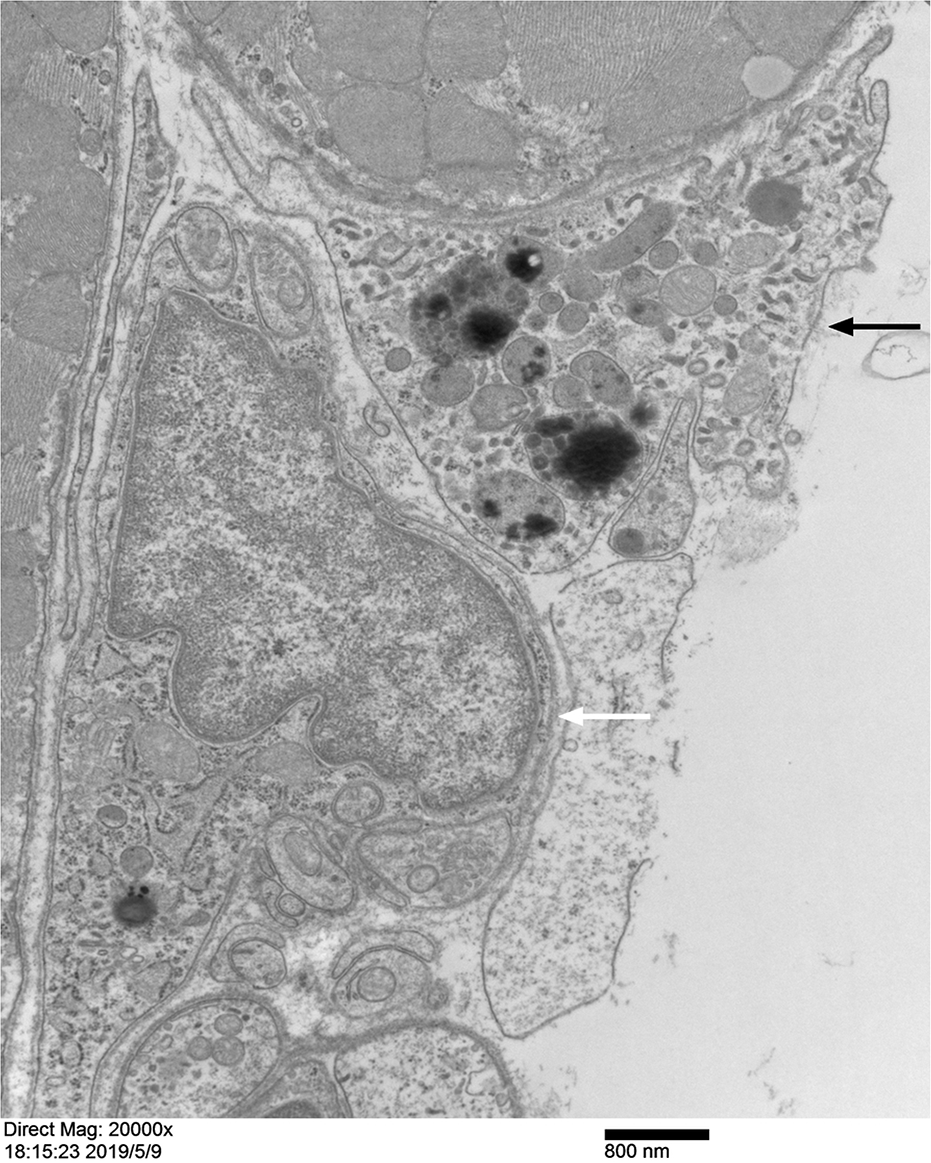
Myocardial tissue around extracellular space in mouse cardiac myocardium. In the myocardial tissue around the extracellular space, a layer of cells is often visible on the surface of the myocardial fibers. This layer of cells can have different cellular characteristics, such as fibroblastic type (shown by the white arrow) or phagocytic type (shown by the black arrow). In addition to rich organelles, the cytoplasm of these cells will contain some lipofuscin-like mass or granules with high or medium electron density. Most of the lipofuscin granules are found within the phagocytic cell. (×20000, scale plate: 800 nm)

In addition, they also contain a large number of lipofuscin granules or clumps of varying sizes and shapes. The elongated cytoplasmic ends of the fibroblasts contain the lipofuscin granules or clumps, which can adhere to the capillaries and fall off. The structural density of the shed cytoplasm at the end of fibroblasts is higher, with higher electron density, and often has structural characteristics that are similar to lipofuscin (Fig 6, 7, 8, and S1, S2 Figs). The basement membrane outside the capillary vessel wall, which is in close contact with the cytoplasm of fibroblasts, often disappears, such that the shed cytoplasm can directly adhere to and fuse with the endothelial cells.

**Fig 8.**
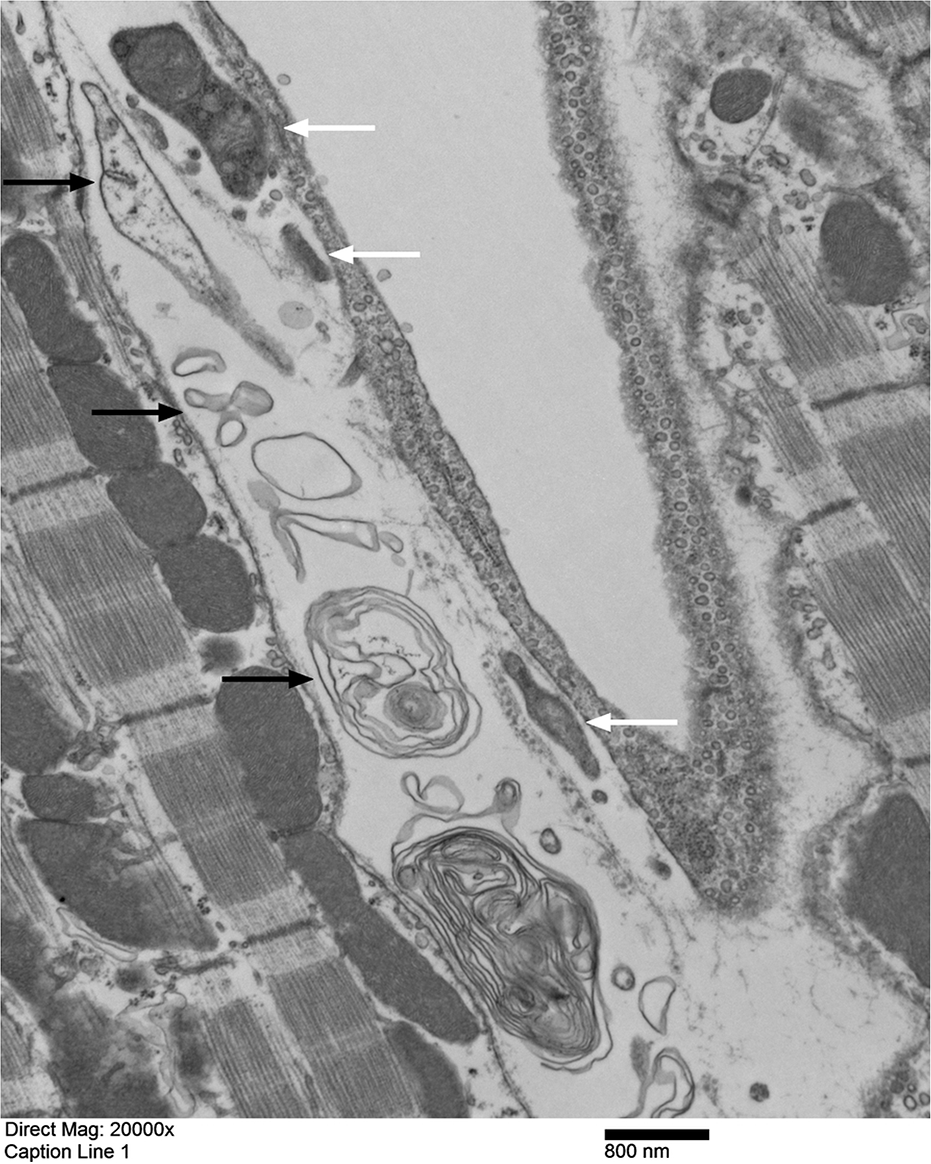
Capillary and perivascular tissue between myocardial fibers in mouse heart. In the interstitial space between the capillary and myocardial fibers, a large amount of messy substance can be seen, some are solid cluster-like structures (shown with white arrows), and some are “membrane-like garbage” (shown with black arrows). The solid clusters have high electron density, an uneven internal structure, and are composed of amorphous substances, which are most likely the cytoplasmic fragments of a fibroblast. The “membrane-like garbage” has a medium electron density, and can form a single layer or multiple layers with irregular structure in size, density and morphology, and partly form lamellar myelin figure structures. The sac-like structures made of these “membrane-like garbage” are all far away from the capillary and close to the myocardial fiber. (×20000, scale plate: 800 nm)

Macrophages are primarily oblong in shape and possess rough surface with several wrinkles or protrusions of different sizes, visibly exhibiting swallowing, phagocytosing and encapsulation of large granules. The internal structure of the cytoplasm is relatively loose and contains abundant vesicles, phagocytosed particles, and higher number of lipid droplet-like lipofuscin granules. These lipofuscin granules vary widely in size, internal structure, and electron density (Fig 5, 7, 9).

**Fig 9.**
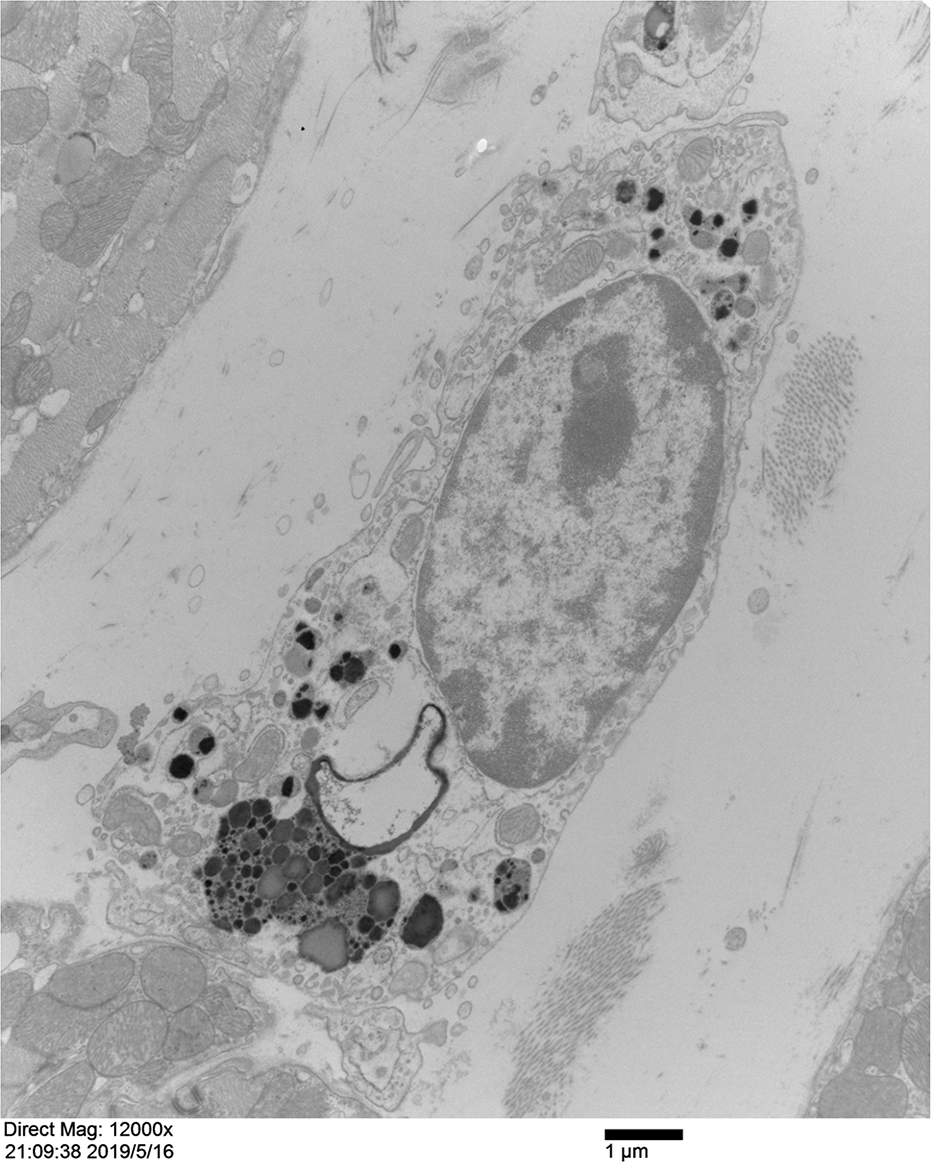
A lipofuscin-containing phagocytic cell in the extracellular space of mouse cardiac myocardium. The image shows a lipofuscin-containing phagocyte in the myocardial space with the morphological characteristics of macrophages. The cell surface is not smooth and has many wrinkles or protrusions of various sizes. There are visible signs of wrapping and swallowing. The cell contains multiple lipofuscin particles and lumps of various sizes, which have higher electron density. Parts of the lipofuscin granules have uneven internal structures with a large difference in electron density. Some lipofuscin granules have lipid droplet-like morphological characteristics. (×10000, scale plate: 1 μm)

Macrophages can either be closely attached to the capillaries through their cell body directly or through the cytoplasm that is elongated and shed. The density of the cytoplasmic fragments shed from the cytoplasmic end of the macrophages is relatively low, and the interior mainly consists of sparse fine particles, vesicles, and lipofuscin granules. The basement membrane outside the capillaries where the macrophages are in close contact with the capillary wall is incomplete or even absent (Fig 5, 10-12). No macrophages were observed in and out of the capillaries or lymphatic vessels.

**Fig 10.**
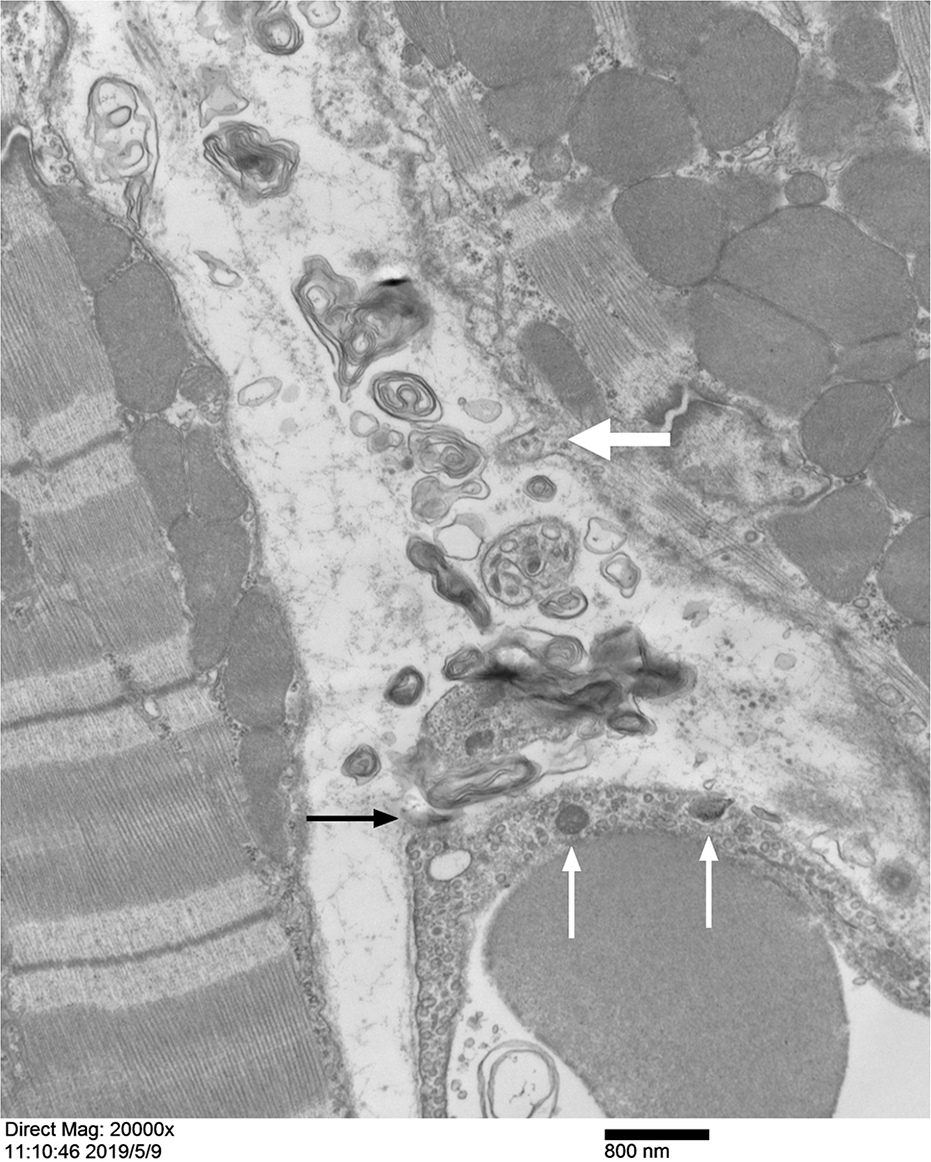
Myocardial interstitium and capillary in mouse heart. A large number of irregular sacs composed of “membrane-like garbage” can be seen in the myocardial interstitium, which have medium electron density and constitute single or multi-layer irregular sac-like structures with large differences in size, density, and morphology. Some membranous substances are densely formed into lamellar myelin figure structures. Some myelin figure structures are close to the capillary wall, there is no basement membrane with the endothelial cell, and even adheres to the endothelial cell (shown by a black arrow). Similar “membrane-like garbage” can be seen in the capillary cavity. On upper part of the capillary wall, oriented toward the membranous myelinfigure structures gathered, there is no basement membrane outside the endothelium. In the endothelium here, two high-density lipofuscin-like granules (shown by thin white arrows) can be seen, and the granule on the right seems to be partially exposed outside the endothelium. In the middle of the surface of the myocardial fiber in the upper right, the surface sarcolemma protrudes into the stroma, forming a rod-like protrusion (shown by a thick white arrow) (×20000, scale plate: 800 nm)

### 2.3. Matrix in myocardial interstitium

In the interstitial matrix of the myocardial space, a large number of non-cellular structured membrane-like structures can be seen, which can be of different sizes and electron density. They form loose, irregular sac-like structures with single or several layers, or dense myelin figures with multiple layers, creating complex structures that greatly vary in size, density, morphology and number of membrane layers (Fig 6, 8, 10-14). We termed these complex membrane-like structures as “membrane-like garbage”. These “membrane-like garbage” are widely and irregularly distributed in the matrix of the myocardial interstitium, either sparsely or densely.

In the interstitium, some free lipofuscin particles can also be seen that vary widely in size. Some are in the form of small granules, while others form large masses, which might be either coated or uncoated at the periphery (Fig 14). Large clumps formed by the aggregation of several lipofuscin granules are sometimes seen (Fig 15). Some lipofuscin granules have low density and a relatively loose structure. The shape is similar to the high-density “membrane-like garbage,” and there is no obvious morphological boundary as that seen in the “membrane garbage,” which is denser in structure (Fig 10-14).

### 2.4. Capillaries in myocardial interstitium

There are many capillaries in the myocardial interstitium, which are continuous in nature. These capillaries can be roughly divided into two types: high density and low density endothelial cell types.

High-density endothelial cell-type capillary vessels have thick walls and high cytoplasmic density. Endothelial cells can form tight junctions. In addition to the scattered mitochondria inside the cytoplasm of the endothelial cells, they contain abundant pinocytotic vesicles and exocytosis vesicles. These vesicles are often highly dense, indicating that active transport occurs through their walls. High electron density granules and fine particles are found abundantly in the cytoplasm (Fig 2, 3, 5, 8, 10-12, 16, and S1 Fig).

The walls of low-density endothelial cell-type capillaries are thinner with relatively less dense cytoplasm. The number of vesicles is low, and few large vesicles and slender membrane tubules can also be seen. The cytoplasm contains many small particles with medium to high electron density, which can adhere to the membrane tube, with morphology similar to the rough endoplasmic reticulum (Fig 14, 16, and S2 Fig).

Morphologically, these two types of capillaries do not have clear boundaries, and the morphology of some vessels is somewhere in between. The endothelial cell membrane on the lumen side of most capillaries has small conical or tiny finger-like protrusions that are of different lengths and can be extruded into the lumen (Fig 2, 14, 16, and S1, S2 Figs). The cytoplasms of some endothelial cells are embedded with lipofuscin-like granules of different sizes and shapes (Fig 9, 10, 14, 16, and S2 Fig). Occasionally, these granules are embedded into the endothelial cells of the vessel walls from the exterior of the vessel without any basement membrane (Fig 10, 11, and S2 Fig). It can also be seen that the lipofuscin granules in the myelin figure are extruded by the endothelial cells into the lumen from the walls of the vessels (Fig 17).

**Fig 11.**
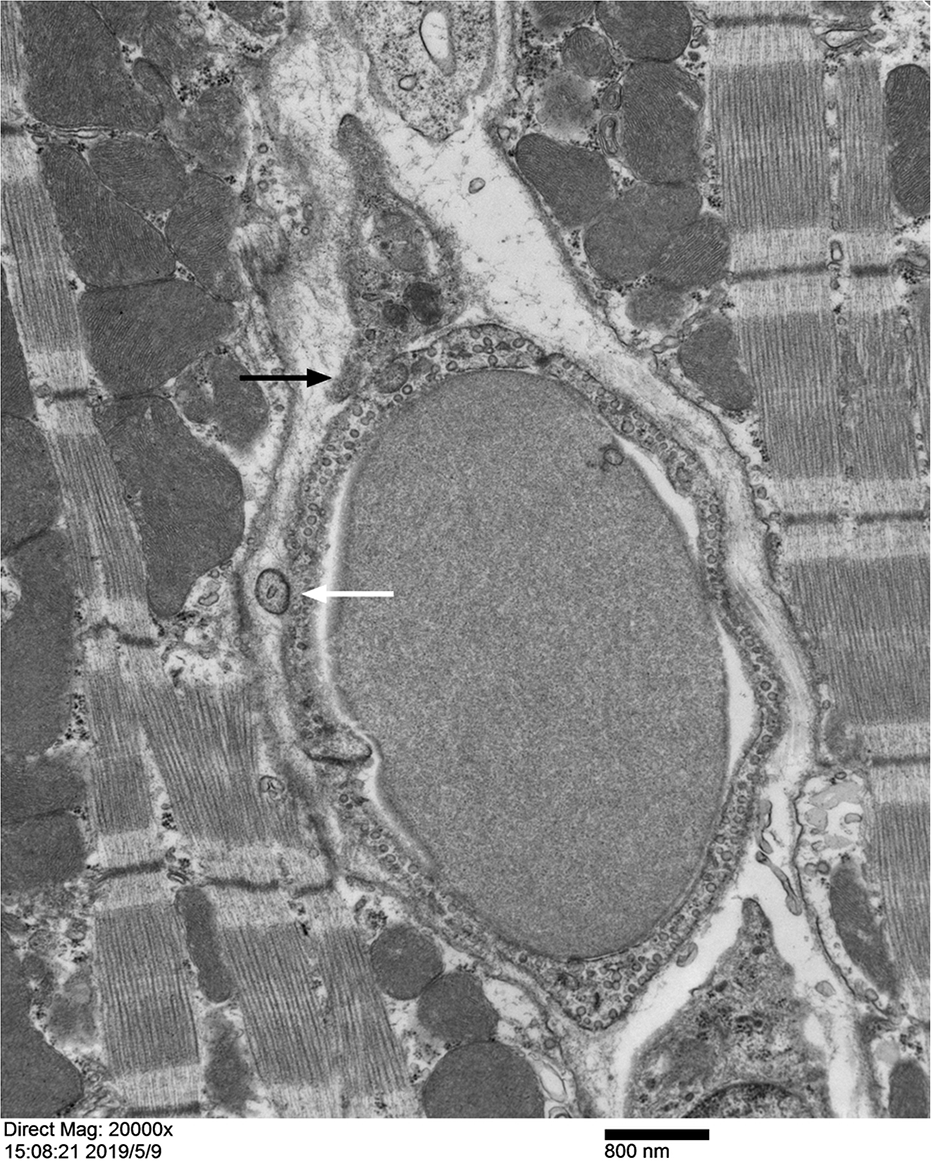
Capillary and surrounding tissue in myocardial interstitium of mouse heart. The central part of the picture is a complete capillary, and the lumen is filled with red blood cells. The vessel wall is surrounded by two endothelial cells, and at the junction of two endothelial cells, tight junctions are formed. In the cytoplasm of endothelial cells of the wall, vesicles are extremely abundant. Most of the endothelium does not have a basement membrane. The endothelium is directly exposed to the interstitial matrix of the myocardium, and is in direct contact with fibers, vesicles, and even myocardial fibers around the capillaries. Slightly above the middle of the left wall of the capillary, a small vesicle can be seen embedded in the endothelium of the capillary wall (shown by a white arrow). In the interstitium above the capillary, a triangular cytoplasmic structure containing several lipofuscin granules can be seen, which is close to the wall of the vessel, and it should be part of the cytoplasmic structure of the lipofuscin-loaded cell in the myocardial interstitial. The lower left side of the cytoplasmic structure is closely attached to the capillary endothelium, and the two parts of the structures seem to be fused (shown by a black arrow). (×20000, scale plate: 800 nm)

“Membrane-like garbage” similar to that found in the myocardial stroma were seen within the lumen of some of the capillaries, which forms sac-like structures formed in the capillary cavity that is more loose and fragmented than the membrane-like garbage formed in the myocardial matrix. The amount of membrane-like garbage in the lumen varies between different blood vessels, and sometimes, there are very obvious differences between the two accompanying blood vessels (Fig 2, 3, 5, 8, 10-12, 14, 16, and S1, S2 Figs). In general, the lumens of the low-density endothelial-type capillaries contain significantly higher amount of membrane-like garbage than the high-density endothelial-type capillaries. This might be because the high-density endothelial-type capillaries are arterial-end capillaries, while low-density endothelial-type capillaries are venous-end capillaries. The blood flowing in the capillaries at the venous end collects more lipofuscin-like substances that are discharged from the capillary walls.

**Fig 12.**
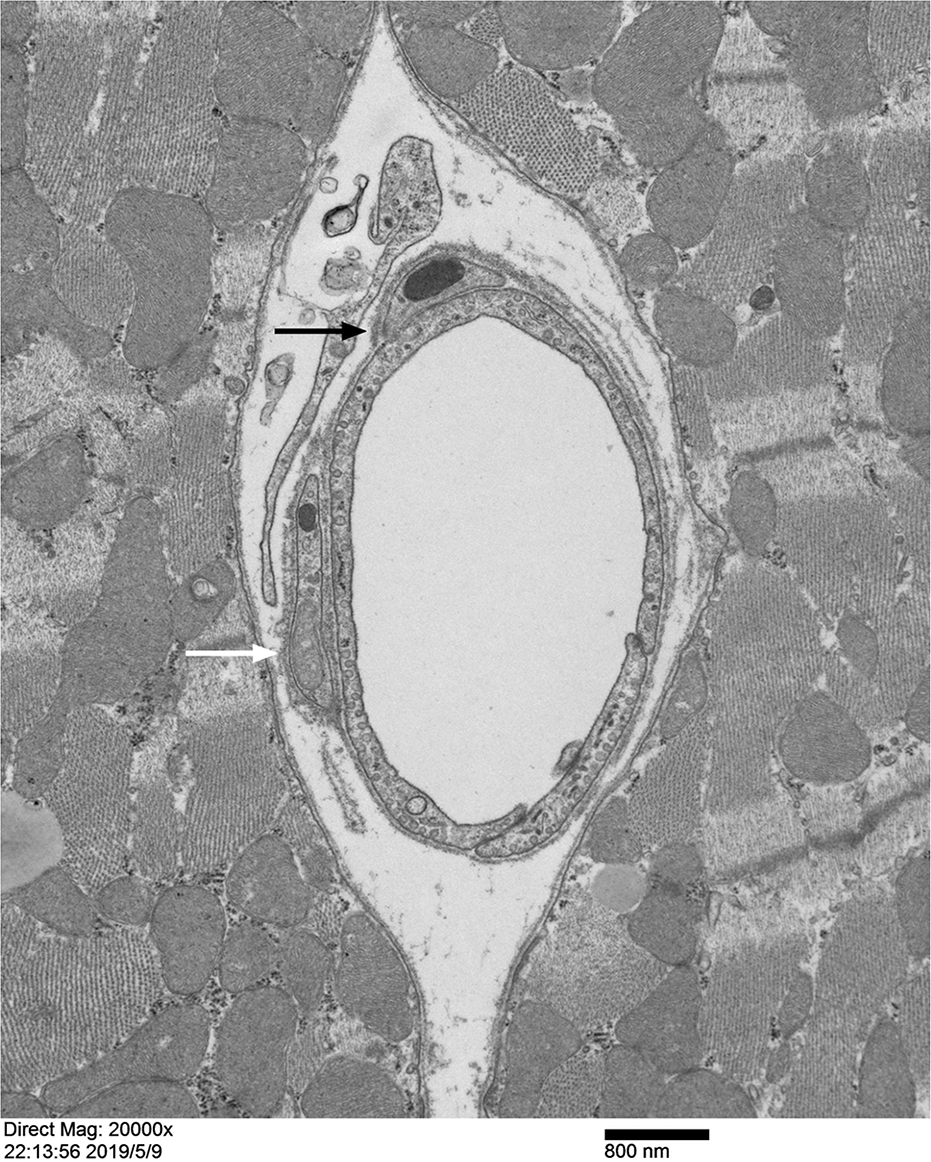
Capillary and outer structure of the vessel wall in the mouse heart myocardium. In the center of the figure is a complete capillary, the vessel wall is surrounded by two endothelial cells. At the junction of two endothelial cells, tight junctions are formed. The cytoplasmic vesicles of endothelial cells constituting the vessel wall are abundant. There is a continuous and complete basement membrane outside the endothelium of the capillary wall. Outside the endothelial wall of the left and upper capillary, each is covered by one non-endothelial cell structure containing lipofuscin-like matter. There is no basement membrane between the non-endothelial cytoplasmic structure located outside the capillary wall and the endothelial cells, and it is closely attached to the capillary endothelium downward, and the two cell structures seem to be fused (shown by a black arrow). The white arrow shows a mitochondrion inside the non-endothelial cell outside the vessel wall. (×20000, scaleplate 800nm)

**Fig 13.**
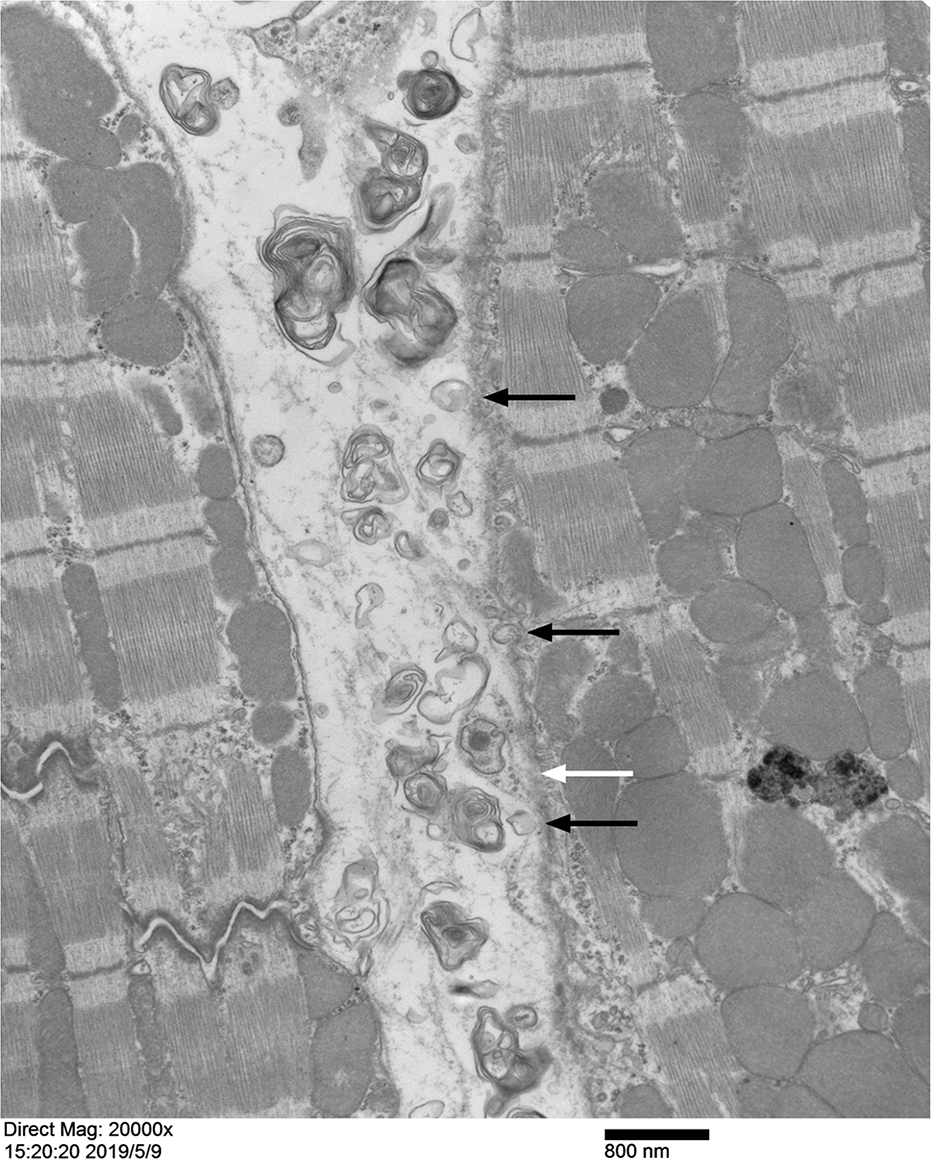
Myocardial fibers and interstitium between myocardial fibers in mouse heart. The image shows the longitudinal section of the myocardial fibers. Wider space is seen between the two myocardial fibers. A large number of randomly distributed “membrane-like garbage” can be seen in the space. These “membrane-like garbage” may form loose, irregular sac-like structures, or densely-formed myelin figure structures. These structures are either loose or dense, with different sizes and shapes, and are scattered in the spaces. The surface of the myocardial fibers on the right side of the space is uneven and the sarcolemma is not very clear. Verruca-like protrusions (shown by white arrow) and several vesicles (shown by black arrows) can be seen on the surface of myocardial fibers. The verruca-like protrusions are broad at the top and slender at the base. A high electron density lipofuscin-like pellet can be seen in the center of the top, and the pellet is surrounded by a thick lipid membrane-like structure. A large lipofuscin mass can be seen deep in the right muscle fiber, with high electron density and uneven internal structure. (×20000, scale plate: 800 nm)

**Fig 14.**
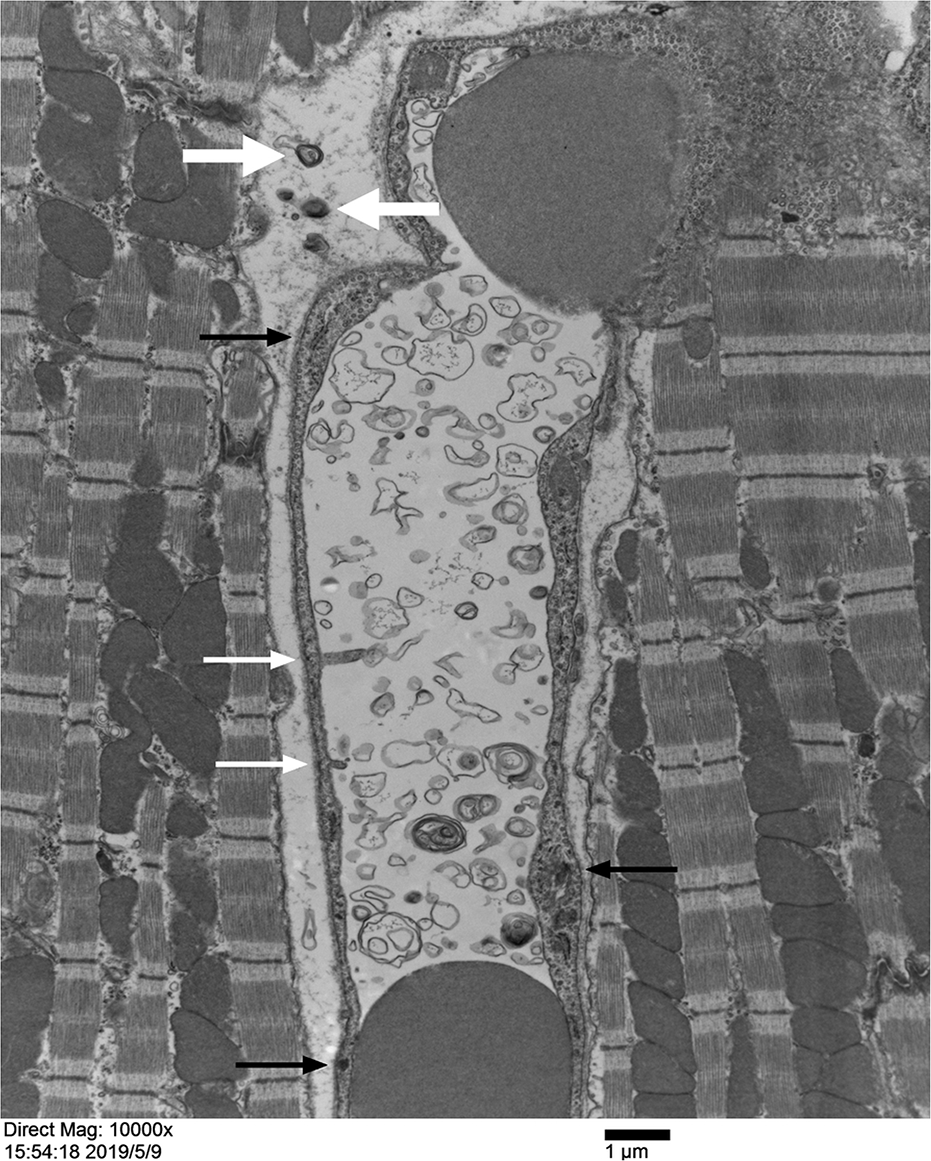
Myocardial fibers and a capillary in the mouse heart myocardium. The center of the figure shows a capillary. The thickness of the endothelial cells constituting the vessel wall varies greatly; the cytoplasm contains abundent vesicles and greater number of pellets or particles with high electron density (shown by black arrows). The cytoplasm of the endothelial cells on the left wall has two mini finger-like protrusions (shown by white thin arrows) that extend to the lumen. In addition to red blood cells, the lumen contains a large number of irregularly shaped “membrane-like garbage” that has large differences in size, density, and shape. Densely structured sacs can form lamellar myelin figure structures. “membrane-like garbage” can also be seen in the stroma outside of the capillary (shown by the white thick arrows). (×10000, scale plate: 1 μm)

**Fig 15.**
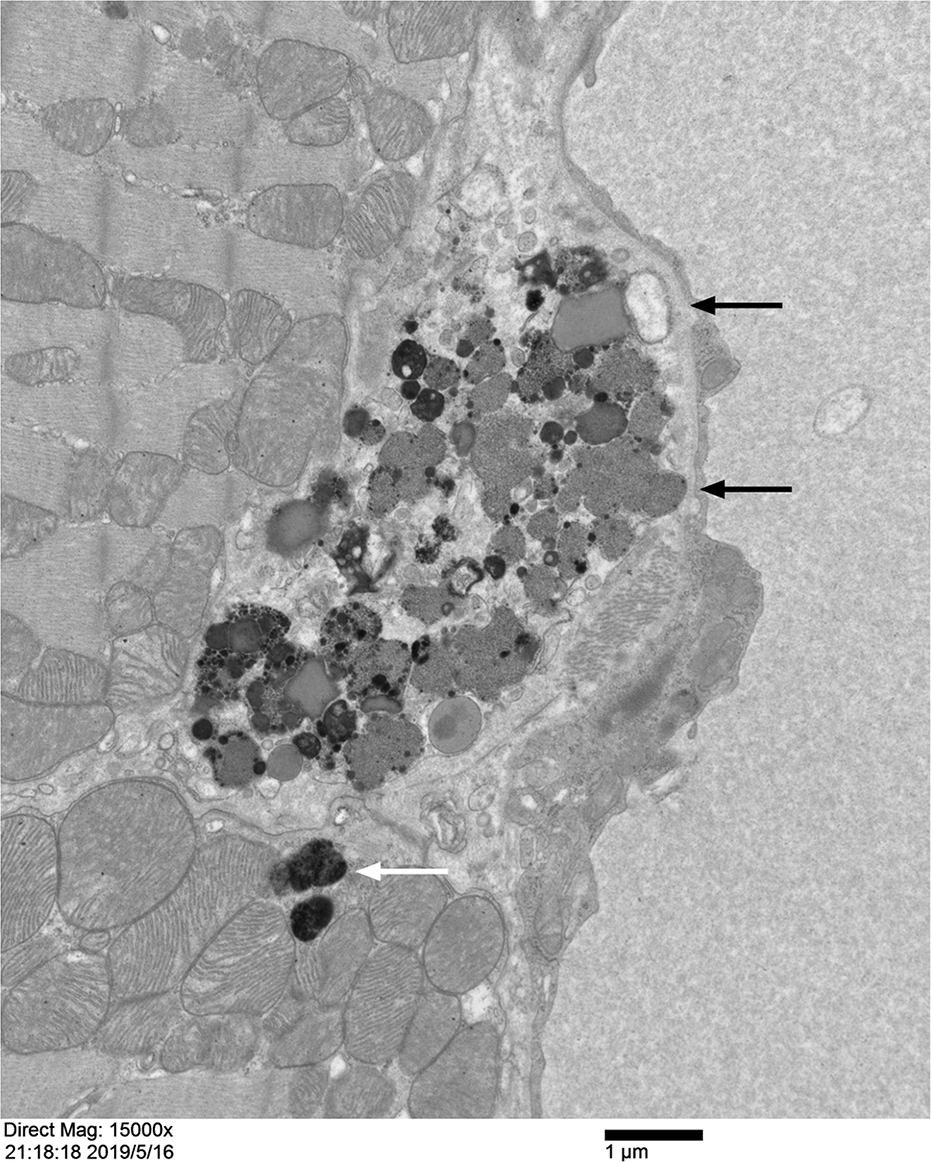
Lipofuscin mass in myocardium of mouse heart. In the interstitium between myocardial fiber and capillary (or lymphatic vessel), there is a large lipofuscin mass. On the periphery of this huge lipofuscinous mass, intermittent membrane wrapping can be seen only in the area below it. Most part of the lipofuscinous mass is directly exposed in the interstitium, directly with the left myocardial fiber and the capillary (or lymphatic vessel) on the right are attached. The basement membrane is not noticeable between the lipofuscin mass and capillary endothelium (shown by black arrows). At the bottom, there are two separate lipofuscin granules, which are located below the sarcolemma of the myocardial fiber (shown by white arrow). (×15000, scale plate: 1 μm)

**Fig 16.**
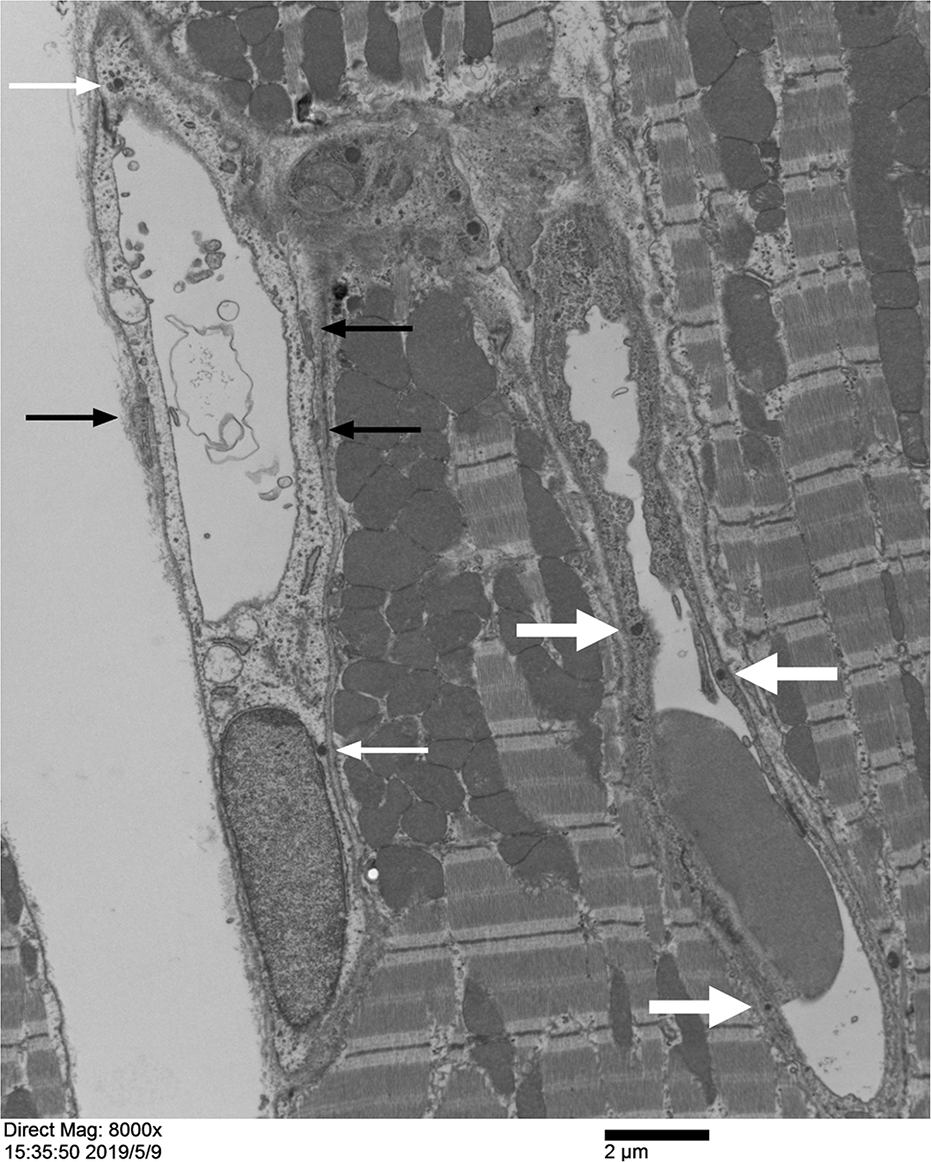
Two capillaries in mouse heart myocardium. The endothelial cytoplasm of the right capillary is rich in vesicles and fine particles with high electron density, which makes the tube wall denser. Several scattered lipofuscin granules (shown by the thick white arrows) can be seen in the cytoplasm. There are fewer intracellular vesicles and high electron density particles in the left capillary endothelium. The structure in the tube wall appears loose, but there are scattered lipofuscin particles (shown by the thin white arrows). On the inner side of the upper part of the left vessel wall of this capillary, the cytoplasm of endothelial cell forms several fine finger-like protrusions that penetrate into the lumen, and fall off. There are several cytoplasmic structures outside the tube wall that are close to the endothelial cells, which shows cytoplasmic fusion (shown by black arrows). (×8000, scale plate: 2 μm)

Capillary vessels are often surrounded by cell bodies or cytoplasmic fragments of different cell types, which can even be directly attached to the vessel wall. The areas where the extravessel cell body or cytoplasmic fragment is in contact with the capillary vessel wall, the basement membrane is often incomplete or even disappears, resulting in direct contact between the extravessel cytoplasm with the endothelial cells, which leads to blurring of the boundaries between them, and even merging. Extravessel cytoplasmic fragments often contain lipofuscin granules (Fig 5, 10-12, and S1, S2 Figs), or are in a highly dense formation (Fig 3, 8, 16, and S1, S2 Figs).

Perivascular cells were not observed around the capillaries. All the cytoplasmic fragments that are outside or surround the capillary walls, and even fuse with endothelial cells, are derived from the extended terminal cytoplasms of the macrophages and fibroblasts, or shed cytoplasmic debris.

## 3. Discussion

Do the heart, brain, and retina have the ability to expel the accumulated lipofuscin? Is it possible to speed up the elimination of lipofuscin in these tissues with drugs such as centrophenoxine and meclofenoxate in order to slow down aging? These are some controversial questions[6, 8, 11, 13, 14, 18–20]. Therefore, this study investigates the elimination of lipofuscin from the heart to obtain potential evidence that lipofuscin is excreted by the tissues. These results might facilitate future research on related interventions.

Observation of myocardial cells revealed that these cells can eliminate lipofuscin. The mechanism of removal of the lipofuscin granules that are located below the sarcolemma might be via the formation of capsule-like protrusions on the sarcolemma and on the sarcoplasm near them, which extend into the interstitial space, detach from the myocardial cells and finally, enter the interstitial space. We did not observe exocytosis of large lipofuscin granules (approximately 1 μm diameter) in the myocardial fibers. This result may indicate that myocardial cells slowly and continuously excrete lipofuscin to the outside in a more subtle way.

Most of the lipofuscin particles entering the interstitium will soon disintegrate, de-aggregate, and diffuse, forming “membrane-like garbage” or mini soluble particles, which are dispersed in the interstitial matrix and diffuse around the blood vessels. These particles can enter the vascular wall of the endothelium through simple diffusion, pinocytosis of capillary endothelium, followed by further diffusion into the bloodstream; or be directly released into the bloodstream by exocytosis. Some mini lipofuscin particles can be adsorbed and fused with other membranous structures (lumen cell membrane, tubules, and vesicles) in the cytoplasm. The vesicles and tubules that fuse with lipofuscin will also merge into the surface membrane of endothelial cell lumen. It cannot be excluded that the mini soluble lipofuscin particles that have diffused into the endothelial cells re-aggregate in the endothelial cytoplasm to form larger lipofuscin granules, which are then “pushed” into the lumen by endothelial cells. This is similar to the phenomenon observed by Joris I *et al*. in human coronary artery where it was seen that endothelial lipofuscin is excreted through enlarged cell processes[15]. A few of the smaller undisintegrated lipofuscin granules might be directly embedded into the capillary endothelium in the form of granules, and then enter the lumen in a “pushed” mode.

Lipofuscin masses in the macrophages might have arisen from three possible ways. First, it might be derived from phagocytosis of intact lipofuscin granules in the matrix. Second, it might be due to micro-engulfing of “membrane-like garbage” along with micro-phagocytosis of fine particles formed by de-aggregation of lipofuscin. These swallowed lipid molecules and phagocytosed particles then aggregate in the cell to form large lipofuscin granules or mass. Third, it might be due to phagocytosis and digestion of other cellular and tissue structural fragments in the cardiac interstitium. The lipofuscin formed through these three methods will eventually aggregate in the macrophages, forming huge lipofuscin masses.

How do macrophages containing large amounts of lipofuscin transfer lipofuscin away from the myocardial tissue? According to our observations and speculations, one of the most significant ways is to transport lipofuscin into the lumen of the blood vessel through fusion with capillary endothelial cells. Thus, the end of the elongated cytoplasm containing lipofuscin granules is close to the capillaries, and it eventually fuses with endothelial cells releasing the lipofuscin granules into the cytoplasm of endothelial cells. The lipofuscin granules entering the vessel wall are transported by the endothelial cells into the lumen in a “pushed” mode, thereby completing the process of transferring lipofuscin from the extravessel stroma into the lumen (Fig 18).

The lipofuscin mass present in the fibroblasts might exist due to two reasons; one is due to the production of its own cell metabolism and the other is due to microphagocytosis, similar to the macrophages. Previous studies have shown that fibroblasts in the myocardial tissue possess a certain phagocytic function[10, 21]. However, in the present study, we did not observe engulfment of lipofuscin granules by the fibroblasts. Fibroblasts also seem to have the ability to excrete lipofuscin and similar to that seen in macrophages, it is excreted into blood vessels by fusion with capillary endothelial cells. Thus, the elongated cytoplasmic ends containing lipofuscin granules is close to the capillaries and fuses with endothelial cells, which releases the lipofuscin material into the cytoplasm of endothelial cells, and finally into their lumen in a “pushed” manner.

As the myocardial tissues we observed are located in the deeper regions of the heart wall, only capillaries that extend between the myocardial fibers, close to the myocardial fibers were visualized, along with a few connective tissues. Therefore, the pericytes or mast cells that were described by other researchers were not observed around the capillaries in our specimens[22].

Comparison of the amount of “membrane-like garbage” in different capillary lumens showed that myocardial tissues “dump” a large amount of “garbage” into the bloodstream during the blood flow, which is mainly derived from lipofuscin produced from myocardial cells, and this dumping is not slow. According to our observations, there are several modes of transport of lipofuscin in the myocardium through the capillary wall (Fig 18). Although there are many modes of transfer, the in-wall and out-wall modes of these transfers are not a fixed combination, and there can be different combinations between different in-wall and out-wall modes. However, from the frequency shown by various transport modes, the cross-wall transport of lipofuscin in myocardial tissue may mainly rely on a small, fine, soluble, continuous transport mode, it is not a one-time, massive, or cell-carrying transfer. This small, fine, soluble, continuous transport pattern is slightly similar to the lipofuscin elimination phenomenon observed in the seminiferous tubules of animal testes[23,24].

Many previous studies have described the presence of lymphatic vessels in the myocardium [25]. However, in the current study, lymphatic vessels with typical structural features were not observed. It is therefore uncertain whether a lipofuscin excretion pathway through the lymphatic vessels exists. This is completely different from the case where the lipofuscin discharged directly into the lymphatic ducts is observed in the testicular seminiferous tubules[23,24].

In this study, we observed that some of the lipofuscin particles were transferred within the myocardial tissue; and across the capillary wall into the blood flow. However, there are still some unanswered questions, such as, is lipofuscin mainly transported in a soluble manner in the myocardium[26]? Is it possible to remove the lipofuscin present in the form of huge clumps? How is it cleared? What roles do fibroblasts play in the elimination of lipofuscin in the myocardium? Do the myocardial tissues from different parts of the mice heart treat lipofuscin in the same way? Further investigations are needed to answer these questions.

**Fig 17.**
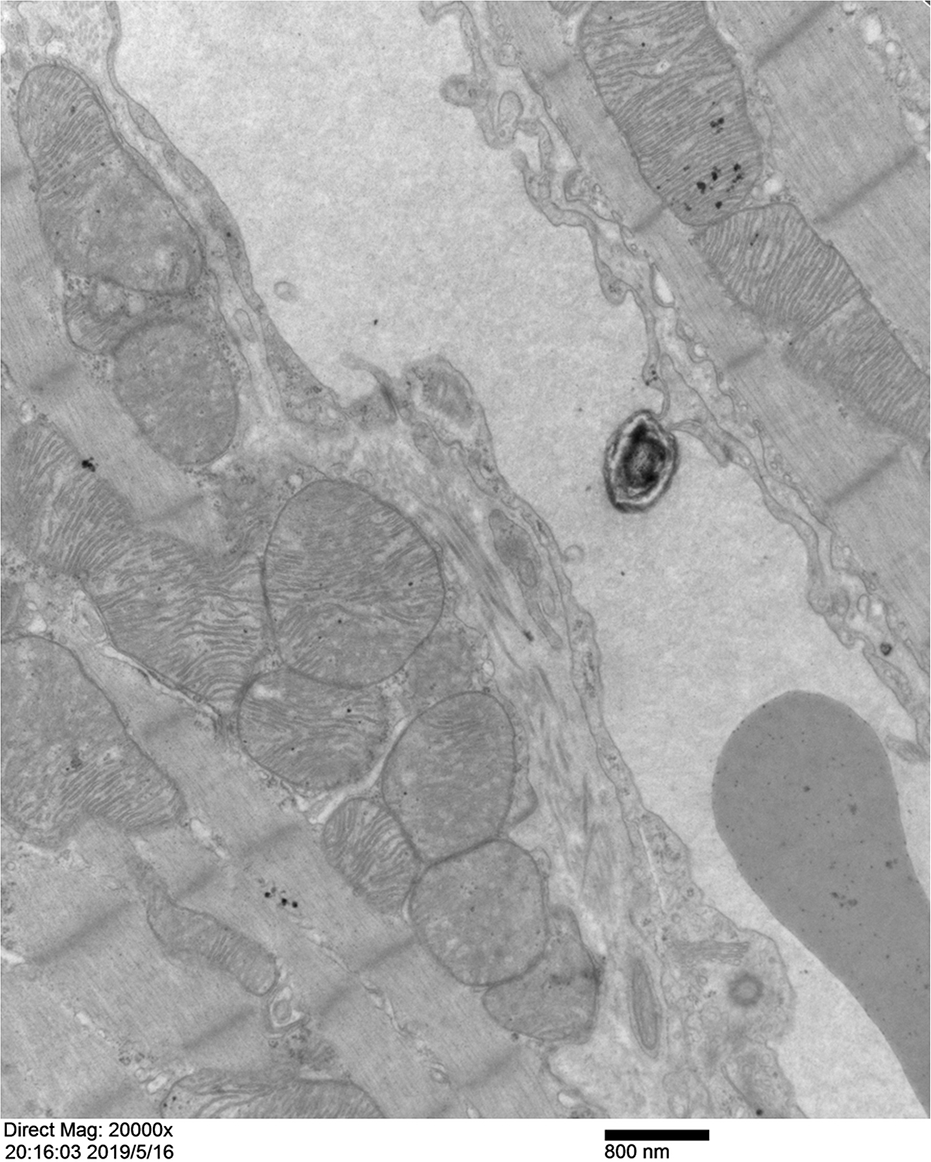
Capillary in the cardiac myocardium of mouse heart. In this picture, a capillary can be seen with a high electron density myelin figure lipofuscin mass protruding from the inside of the vessel wall into the lumen at the center of the right vessel wall. (×20000, scale plate: 800 nm)

**Fig 18.**
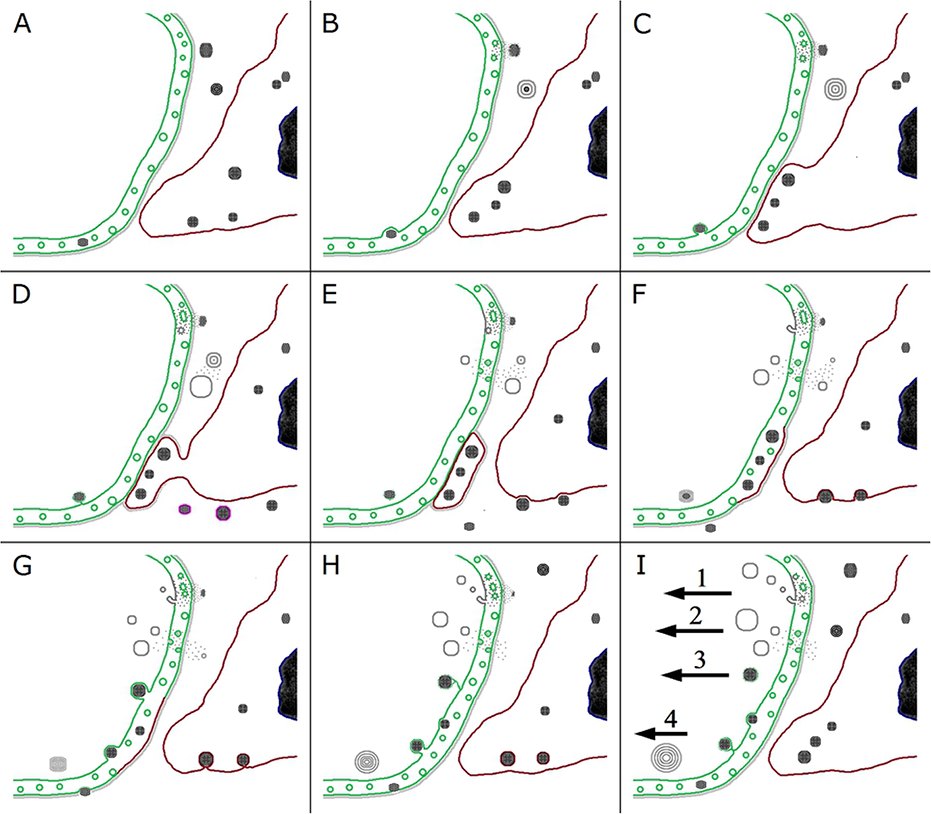
Schematic diagram of the lipofuscin excretion through the capillaries in the myocardial interstitium of the mouse heart. The figure shows four possible pathways through which the lipofuscin found in the myocardial interstitium enters the capillary cavity. Pathway 1: The lipofuscin granules are close to the capillary wall, and the granules gradually disaggregate to form soluble micro-particles. These microparticles enter the endothelium of capillaries by simple diffusion, and then aggregate in the cytoplasm of the endothelial cells to form fine lipofuscin particles. These particles can adhere and subsequently fuse with the membrane-bound vesicles and tubules in the cytoplasm, as well as the cell membrane on the lumen surface. Membrane vesicles and tubules fused with lipofuscin can further merge with the cell membrane on the lumen surface. The cell membrane of the luminal surface with lipofuscin forms fine finger-like protrusions into the cavity and these protrusions continue to elongate and eventually fall off to enter the blood stream. Pathway 2: The lipofuscin granules swell and dissolve, forming a lamellar membrane-like myelin figure structure, and further disaggregate to form soluble microparticles or lipid molecules, which can re-aggregate in the interstitium to form irregular membrane sacs, or they can diffuse directly through the basement membrane. Irregular membrane sacs formed by re-aggregation in the interstitium will soon disaggregate to form soluble microparticles or lipid molecules that can diffuse through the basement membrane and enter the capillary lumen directly through simple diffusion, or they can be swallowed by the endothelium to form pinocytotic vesicles, which are transferred into the cavity. Finally, these micro-particles and lipid molecules are expelled into the lumen by exocytosis where they can re-aggregate. In the circulating plasma, they aggregate again to form membrane vesicles of different sizes and shapes. Pathway 3: Lipofuscin granules in the interstitium enter the fibroblast cells or macrophages, and these cells can extend their lipofuscin-containing cytoplasm to the capillary and closely adhere to the vessel wall. The protruding cytoplasmic portion containing lipofuscin granules can get detached from the original cell body. The basement membrane outside the endothelium of the capillary wall that is attached to the cytoplasm of lipofuscin-containing cell disappears, and the two cells closely adhere. Finally, these cells are fused, and the lipofuscin-containing cell cytoplasm merges with the endothelium. Simultaneously, the lipofuscin granules that are brought into the endothelial cells can be “pushed” into the capillaries. Pathway 4: Lipofuscin granules pass through the basement membrane, directly enter the capillary endothelium, and then are extruded into the lumen of the blood vessel in a “pushed” manner. They finally fall off from the endothelium to enter the blood stream where they rapidly swell and disintegrate to form lamellar myelin figure structures with a relatively loose structure, and further disaggregate to form “membrane-like garbage”.

## Supporting information

**S1 Fig. Capillary and fibroblast in the cardiac myocardium of mouse heart.**

A complete capillary can be seen in the myocardial interstitium. The lumen of the capillary is almost filled with red blood cells, leaving a little space on the top. There are several elongated finger-like protrusions from the endothelial cell. Below the capillary, a fibroblast can be seen, with an elongated protrusion, and it is closely attached to the outer wall of the capillary. In this area, there is no basement membrane notieable, and thus, the end of the protrusion is directly attached to the endothelial cell (shown by black arrows). Above the capillary, there are two small lipofuscin-like granules (shown by white arrows). (×20000, scale plate: 800 nm) (shown by thin white arrows)

**S2 Fig. Structure of a capillary in the myocardium of the mouse heart.**

It can be seen that the capillary wall is thick, structural density of the endothelial cell is low, and the number of vesicles is low. In the cytoplasm of endothelial cell, three medium-high electron density spherical lipofuscin granules (shown by black arrows), few medium-high electron density fine particles, and several medium-high electron density tubules can be seen. The left inner wall of the capillary is not smooth, and the endothelial cell membrane protrudes into the lumen, forming fine finger-like protrusions and vesicles. Membrane sacs and membrane debris-like structures are scattered throughout the capillary cavity. The exterior of the vessel wall on both sides contains cytoplasmic fragments of extramural cells attached to the exterior of the endothelial cell (shown by white arrows). (×20 000, scale plate: 800 nm)

## Authors’ contributions

Lei Wang performed the morphological observation, animal feeding and sample collection. Chang-Yi Xiao organized or participated in the design of the study and performed the morphological observation.

Jia-Hua Li and Gui-Cheng Tang were responsible for the preparatory laboratory work, dissection, and initial preparation and processing of the samples.

Xiao Rong-Shuang provided help in modifying the English translation of the manuscript. All authors read and approved the final manuscript.

